# Large-scale mapping of positional changes of hypoxia-responsive genes upon activation

**DOI:** 10.1101/2021.09.27.462053

**Authors:** Koh Nakayama, Sigal Shachar, Elizabeth H. Finn, Hiroyuki Sato, Akihiro Hirakawa, Tom Misteli

**Author notes:** Corresponding authors: Koh Nakayama, Ph. D., Department of Pharmacology, School of Medicine, Asahikawa Medical University, 2-1-1-1 Midorigaoka higashi, Asahikawa, Hokkaido 078-8510, Japan; Tom Misteli, Ph.D., Cell Biology and Gene Expression Group, Center for Cancer Research, NCI, NIH, Bethesda. **Abbreviations** HIPMap: High-throughput imaging position mapping, HIF: Hypoxia-inducible factor, VEGF: vascular endothelial growth factor, ZEB: zinc finger E-box binding homeobox, HT: HIF-target, CDT: CREB-direct target, ICT: indirect CREB target, NC: non-responsive control.

## Abstract

Chromosome structure and nuclear organization are important factors in the regulation of gene expression. Transcription of a gene is influenced by local and global chromosome features such as condensation status and histone modifications. The relationship between the position of a gene in the cell nucleus and its activity is less clear. Here, we used high-throughput imaging to perform a large-scale analysis of the spatial location of a set of nearly 100 hypoxia-inducible genes to determine whether their location within the nucleus is correlated with their activity state upon stimulation. Radial distance analysis demonstrated that the majority of HIF- and CREB-inducible hypoxia responsive genes are located in the intermediate region of the nucleus. Radial position of numerous responsive genes changed upon hypoxic treatment. Analysis of the relative distances amongst a subset of HIF target gene groups revealed that some gene pairs also altered their relative location to each other upon hypoxic treatment, suggesting higher order chromatin rearrangements. While these changes in location occurred in response to hypoxic activation of the target genes, they did not correlate with the extent of their activation. These results suggest that induction of the hypoxia-responsive gene expression program is accompanied by spatial alterations of the genome, but that radial and relative gene positions are not directly related to gene activity.

## Introduction

All cells in the human body constantly consume oxygen for efficient energy production, and multiple biochemical reactions are mediated by oxygen-dependent enzymes under normal conditions (Losman *et al*., 2020). When cells in an organism are exposed to low oxygen levels, the hypoxic response is triggered (Kaelin and Ratcliffe, 2008). The hypoxic response is an active process that involves the induction of a specific gene expression program that counteracts the potentially harmful effects of hypoxia to cells and tissues. <u>H</u>ypoxia-Inducible <u>F</u>actor, HIF, is a master transcription factor of the hypoxic response and plays a critical role by up-regulating a number of hypoxia-responsive genes (Keith *et al*., 2012). HIF induces a wide range of genes including many involved in vasculo-/angio-genesis, red blood cell production, and regulation of metabolism (Hirota, 2020). HIF-1α expression decreases upon prolonged hypoxic treatment, and CREB and NF-κB become activated during this stage (Nakayama, 2013).

The activity of genes is affected by their local chromatin environment. In particular, there are two major types of chromatin: heterochromatin which has higher density and often contains transcriptionally repressed genome regions (Passarge, 1979) and euchromatin which is more decondensed and contains transcriptionally active genes (Jost *et al*., 2012) as well as poised genes which become transcriptionally active in response to cellular signals. Gene activation often involves changes in chromatin structure, including decondensation of chromatin or association of distal regulatory regions with promoter elements via long-range loop formation (Dekker and Misteli, 2015). The activity of genes has also been linked to their position relative to nuclear structures such as the association of inactive genes with the nuclear periphery (Takizawa *et al*., 2008b; Egecioglu and Brickner, 2011) and of active genes with nuclear splicing speckles (Spector and Lamond, 2011).In addition, it has been suggested that co-regulated genes may cluster around shared transcription sites in the cell nucleus (Osborne *et al*., 2007). However, the relationship of gene location with activity has not been probed systematically in a large set of genes.

The hypoxic response requires induction of multiple genes in a coordinated manner in order to adapt to low oxygen and as such offers an opportunity to map the location of a set of co-regulated genes in response to a specific milieu. In the present study, we have used the cellular response to hypoxia as a model system to probe the relationship between 3D gene position and activity. We performed positional mapping of 104 hypoxia-responsive genes using HIPMap, a High-throughput Imaging Positioning MAPping method based on fluorescence in-situ hybridization (FISH) using barcoded probes, which allows determination of 3D locations and distances at large scale (Shachar *et al*., 2015a, Shachar et al., 2015b). We find evidence for large-scale re-organization of the genome in response to hypoxia but do not detect any correlation between radial or relative spatial position and gene activity nor evidence for spatial clustering of co-regulated genes.

## Results

### Hypoxic treatment in high-throughput format

To probe the relationship between gene position and gene activity we used high-throughput FISH to comprehensively map the nuclear position of a set of 104 hypoxia-inducible genes in human MDA-MB231 breast cancer cells which are sensitive to oxygen levels (Nakayama, 2013). Cell culture conditions and hypoxic treatment of 1% oxygen were optimized for use in the 384-well plate format using a customized hypoxia chamber and standard fixation and imaging methods were applied (Figure 1A; see Materials and Methods). Immunocytochemistry demonstrated the expected induction by 9.3±2.9-fold (p=3.52 x 10^-26^) in expression of HIF-1α after 24 h of hypoxic treatment compared to cells grown at normal oxygen levels (Figure 1B and Supplemental Figure 1). Similarly, phosphorylation of CREB, a factor activated under prolonged hypoxia (Nakayama, 2013), was observed at 48 h of hypoxic treatment (Figure 1B). HIF- 1α staining was positive in all wells examined, indicating the absence of position and edge effects within the plate (Supplemental Figure 1). Induction of hypoxia was also confirmed by lactate assays which showed higher lactate level in hypoxic samples compared to the normoxic samples (Figure 1C; norm 17.3±0.88 μM, hypo 48.6±3.00 μM, p=3.88 x 10^-7^). Since the nuclear size and shape also affect gene position, we compared nuclear morphology by calculating nuclear area and nuclear roundness. Normoxic and hypoxic treated cells both had a nuclear area of 160 μm^2^ on average, and similar nuclear roundness, suggesting that nuclear shape and area do not change upon hypoxic treatment (Figure 1D).

**Figure 1:**
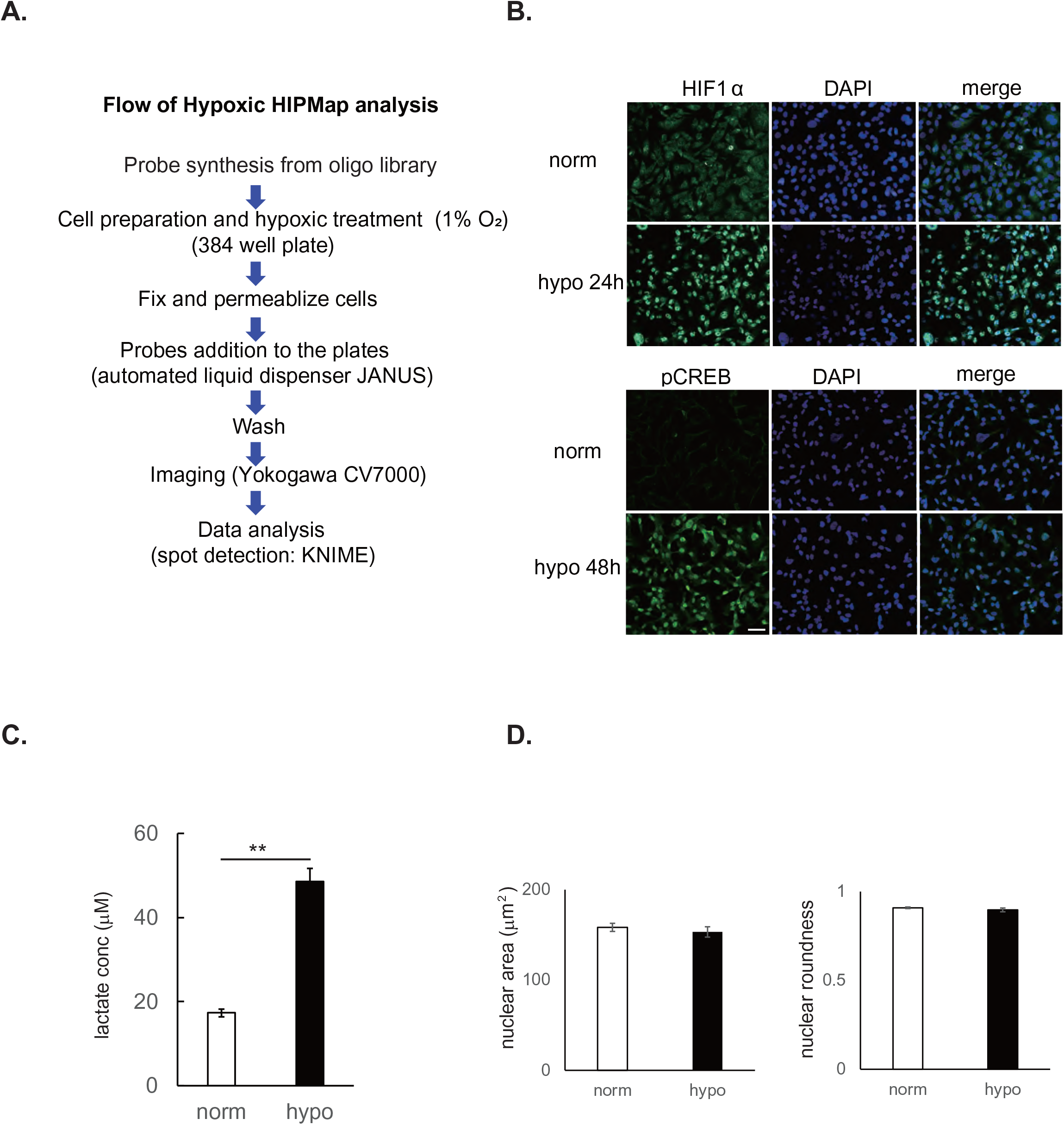
HIPMap analysis under hypoxic condition. **A.** Schematic overview of hypoxic HIPMap analysis. **B.** MB231 cells plated in 384-well plate format were treated with normoxia (21% O2) or hypoxia (1% O2) for the indicated time. After fixation, cells were stained with anti-HIF1α or anti-phospho CREB (pCREB) antibody. Nuclei were stained with DAPI. Scale bar: 50 μM. **C.** Lactate assay of hypoxic-treated MB231 cells in 384-well plates. Cells were maintained under hypoxic condition (1 % O2) for 48 h. Cell culture medium was collected and subjected to lactate assay. Values represent averages of the experiments (n=6), error bars indicate SD. Significance was analyzed by t-test (** p<0.02). **D.** Nuclear area and nuclear roundness were calculated in normoxic and hypoxic treated cells using Columbus image analysis software (Perkin Elmer inc.,). Values represent averages of experiments (n=6), error bars indicate SD. The difference between normoxic and hypoxic samples was not significant.

### Positional mapping of hypoxia-inducible genes in the nucleus

With this experimental setting, we performed a high-throughput FISH analysis by HIPMap (Shachar *et al*., 2015). 104 hypoxia-responsive genes were selected based on their dependence on the two major hypoxia transcription factors HIF and CREB. HIF-target genes were identified based on previous expression studies (Wenger *et al*., 2005), and putative CREB target genes were identified from RNA-seq analysis comparing WT and CREB-knockdown MB231 cells (Kikuchi *et al*., 2016). Target genes were grouped in three major groups: 35 known direct HIF targets (HT) which contain HIF binding sites; 3 direct CREB targets (CDT), which contain CRE motifs in their promoter region; and 64 indirect CREB targets (ICT), which were decreased in CREB-KD cells but do not have CREs (Kikuchi et al. 2016). 2 genes which did not show any significant difference between normoxic and hypoxic conditions in the RNA-seq analysis were used as controls. Oligo probe-based FISH accurately detected typical gene signals in the nucleus of MB231 breast cancer cells which are mostly triploid (Figure 2A).

**Figure 2:**
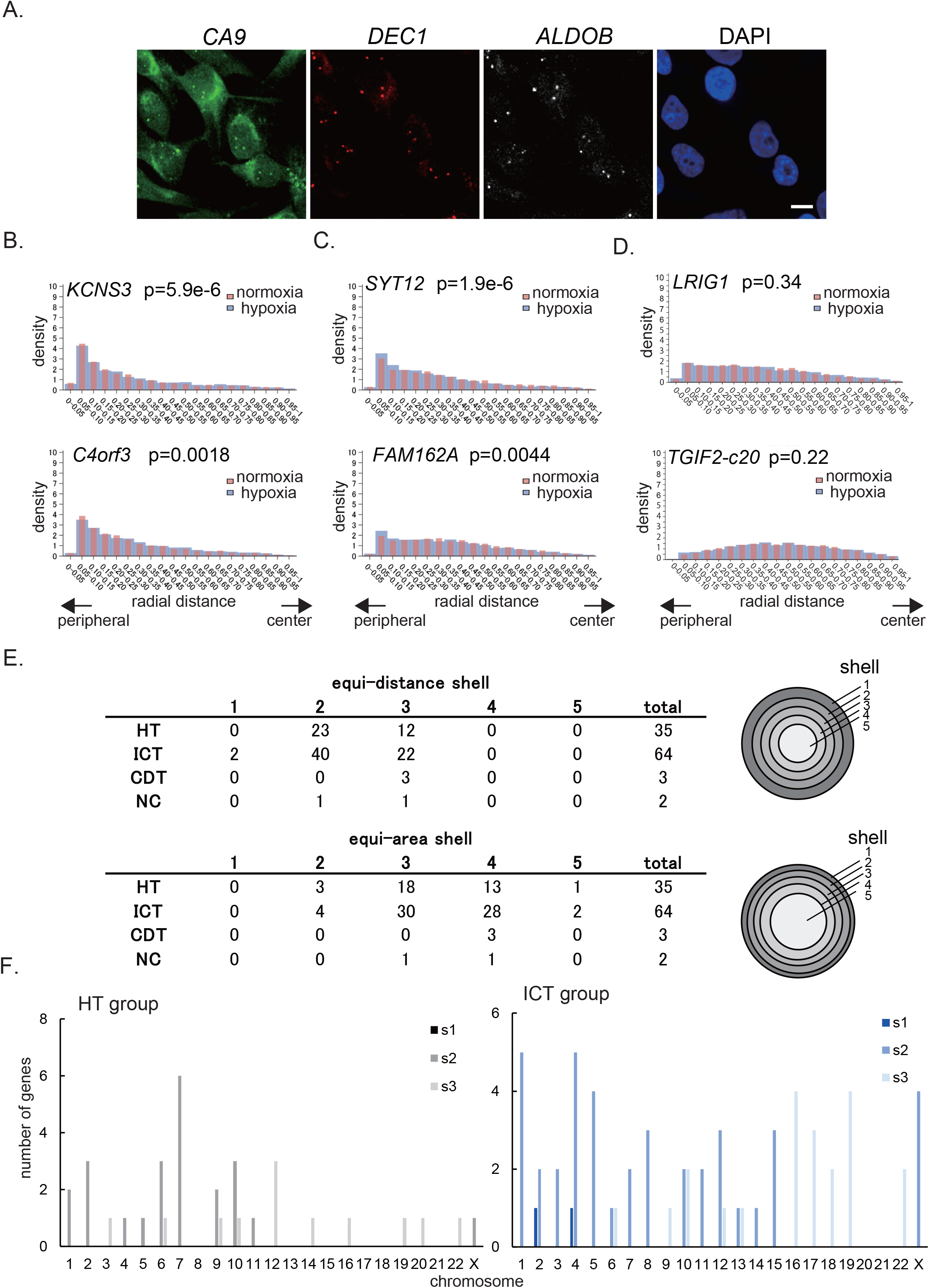
Radial Positions of HIF and CREB target genes under hypoxia. **A.** Representative images of Oligo paint-based FISH. Three hypoxia-inducible genes on chromosome 9; *CA9* (green), *DEC1* (red), *ALDOB* (white). Nuclei were stained with DAPI (blue). Scale bar: 10 μM. **B., C., D.** Radial position of representative genes. Radial positions of genes comparing normoxia and hypoxia were plotted in a histogram (red bar; normoxia, blue bar; hypoxia). Plots represent the data from FISH spots. B.: genes shifted toward center of the nucleus upon hypoxia. C.: genes shifted toward peripheral of the nucleus upon hypoxia. D: control genes. **E.** Radial positions of hypoxic genes. Distribution of the genes examined were shown in five concentric equi-distance and equi-area nuclear shells. Shell 1 is the most peripheral region of the nucleus, and shell 5 is the most central region. HT: HIF-target group, ICT: Indirect-CREB target group, CDT: CREB direct target group, NC: non-responsive control. **F.** Relationship between radial position and chromosome location. Genes of the HT and ICT group are classified based on their chromosome location and plotted in relation to the radial position (equi-distance nuclear shell number(s)1, 2 or 3). Gene number was counted in the integrated data of different experiments. Black/grey: HT group, blue/light blue: ICT group.

To determine the nuclear position of these genes, we calculated the nuclear radial position of each gene at the allele level, normalizing each measurement to nuclear size as previously described (Gudla *et al*., 2008). For each experiment, done with at least two biological replicates per gene, we compared normoxic and hypoxic conditions, and calculated 1) mean radial position in each condition, 2) the variance in radial position in each condition, and 3) the p-value to compare induced vs uninduced position using the Wilcoxon rank sum test. Genes were filtered for genes which showed a consistent change across biological replicates.

To determine the degree of change in gene position, we used the inverse variance-weighted method to calculate the weighted ratio of mean radial distance for each experiment and corresponding 95% confidence intervals (Friedrich et al. 2011). For comparisons between hypoxic to normoxic conditions, multiple p-values generated by the Wilcoxon rank sum test on independent experiments were combined using Fisher’s method (Fisher 1932).

Amongst 104 genes (Supplemental Table 1) the radial distance distributions showed that each gene has a unique, non-random positioning pattern within the nucleus of MB231 cells (Figure 2B-D). Most genes showed a distinct peak at a particular distance from the nuclear membrane, rather than a broad distribution across the interior-exterior axis, suggesting that they have a preferred radial position in the nucleus as previously observed (Shachar et al., 2015a, Meaburn et al., 2009). No characteristic distribution profiles were associated with HT or ICT group genes. Typically, about half of the genes were located in the most peripheral region and the other half were located in the intermediate region, with no genes enriched in the nuclear interior (Figure 2B-C). Similar distribution patterns were also observed for the control gene group (Figure 2D). We further grouped the genes by equi-distant or equi-area shells in the nucleus, based on the median of radial distances. We calculated the median radial distance for each gene to determine the corresponding shell location. All genes analyzed were found in the peripheral shells 2 or 3, regardless of the gene group in the equi- distance analysis, whereas the distribution showed greater variation in the equi-area analysis which ranged from shell 2 to 5 (Figure 2E). As expected, genes on the same chromosome were generally located in the same shell, which confirms that radial position indicates the relative chromosome location in the nucleus (Figure 2F, Supplemental Figure 2 and Supplemental Table 2).

To ask whether activation of hypoxia-responsive genes leads to a change in their nuclear location, we compared the radial position of 92 genes under normoxia and hypoxia conditions (Supplemental Table 1, bold genes). Amongst this set, the radial distribution of 21 genes differed statistically between hypoxia and normoxia (Figure 2B, C and Supplemental Table 3; P <0.05). Out of the 21 genes, 2 were from the HT group, 1 was from the CDT group, and the remaining 18 were from the ICT group. Enrichment of CREB target genes amongst repositioning genes was statistically significant by hypergeometric analysis (p=0.031). Neither of the control genes changed their position in this analysis. 15 genes moved towards the interior of the nucleus (hypoxia / normoxia mean > 1), whereas 6 genes moved toward the periphery of the nucleus upon hypoxic treatment (hypoxia / normoxia mean < 1; Supplemental Table 3). Expression of 11 genes out of 15 was up-regulated in the internally-shifted group, while 5 genes out of 6 were up-regulated in the periphery-shifted group. The largest changes in radial positioning were observed for a potassium voltage-gated channel modifier, *KCNS3*, whose mean distribution shifted toward the interior of nuclei, and synaptotagmin *SYT12* which moved toward the periphery of nuclei. These changes were accompanied by changes in expression level of *KCNS3* which decreased 1.12 fold, and *SYT12* which increased 1.23 fold (for the profile of all 21 genes; see Supplemental Table 3). The observed changes in radial position were unrelated to the chromosomal location of these genes. For example, there were 8 genes on chromosome 7 examined, and only one gene significantly changed its radial position (Supplemental Table 2 and 3). This result indicates that the changes in gene position are local events occurring at the level of individual loci and not repositioning of entire chromosomes.

### Relative distances of hypoxia-inducible genes under hypoxic condition

We next determined the position of HIF-responsive genes relative to each other by measuring the pairwise distances between FISH signals for 159 pairs of genes located on different chromosomes (Supplemental Table 4). In this analysis, we focused on the HT group and analyzed the shortest pair-pair distances of each allele (Figure 3A). Given that these genes had a relatively constant radial position, we were interested in their relative position. Data analysis was performed in the same way as the radial distance analysis by calculating the weighted ratio of means of relative distances (hypoxia / normoxia) and corresponding 95% confidence intervals (Friedrich et al. 2011), and the combined p-value on the Wilcoxon rank sum test using Fisher’s method (Fisher 1932). Out of the 159 pairs, 74 pairs showed statistically significant and reproducible differences in the relative distance between normoxia and hypoxia (Figure 3B, C; Supplementary Table 5; P < 0.05). Based on the ratios of means of the relative distance between normoxia and hypoxia, 30 pairs became more distal (green), whereas 44 pairs became more proximal (red) (Figure 3D and Supplemental Table 5).

**Figure 3:**
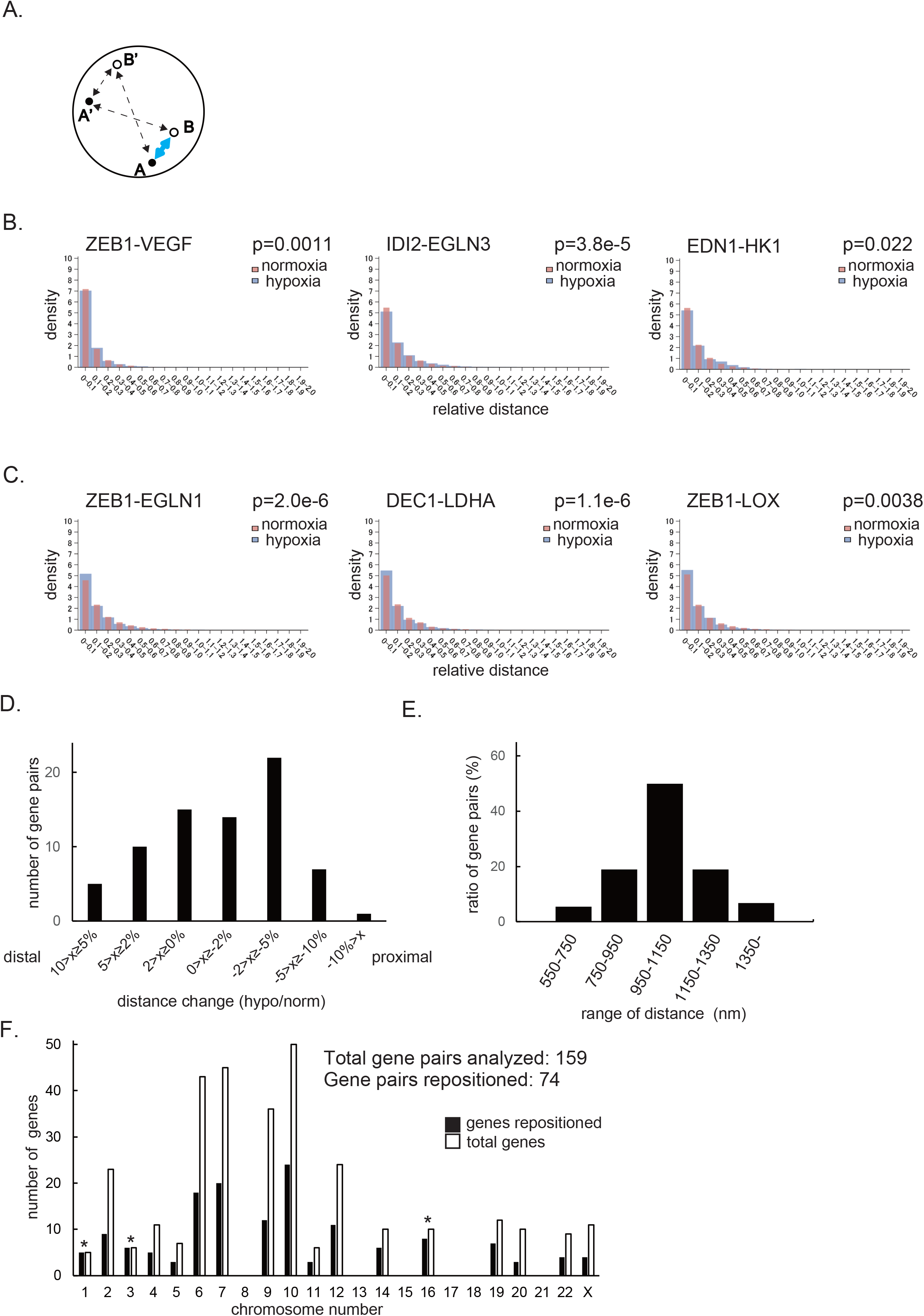
Relative distances of HIF target genes under hypoxic condition. **A.** Relative distances of hypoxia-inducible genes in the nucleus. Minimal distance of hypoxia-inducible genes (A-B) were calculated under normoxic and hypoxic conditions (blue solid arrow). **B. , C.** Distribution of gene distances comparing normoxic and hypoxic conditions. Changes in the distance of gene pairs was measured and plotted. Distance of gene pairs which became more distal (B.) and more proximal (C.) upon hypoxia are shown (red bar; normoxia, blue bar; hypoxia). **D.** Number of gene pairs which became distal or proximal. Gene pairs which changed the distance at most in either direction were counted in the integrated data of different experiments and plotted. X axis indicate the degree of distance change (hypo/norm). **E.** Distance distribution of gene pairs. Mean distance of gene pairs were calculated for the 74 genes which showed significant differences under hypoxic conditions. **F.** Genes with frequent movement based on chromosome location. Genes which changed position were grouped based on the chromosome. Numbers of total genes analyzed and repositioned genes on each chromosome are shown. * :p<0.05.

The mean pair-to pair distances under hypoxic condition ranged from 586 to 1697 nm, and mean pair distances below 500 nm did not exist (Figure 3E). Colocalization within one pixel ranged from 21.8% to 49.5% (average 31.3%) and was strongly correlated with mean distance. Although there was no clear indication of gene clustering, about 60% of the gene pairs analyzed moved apart upon hypoxia (P < 0.05). Pairs of genes which showed altered distances were enriched on chromosomes 1, 3 and 16 (p=0.02, 0.009, and 0.029, respectively). Among the 74 gene pairs (148 genes), 5 genes (out of 5) were on chromosome 1, 6 genes (out of 6) were on chromosome 6, and 8 genes (out of 10) were on chromosome 16 (Figure 3F). These results demonstrate the absence of clustering of co-regulated genes but indicate that some chromosomes regions undergo local changes in response to hypoxic stimulation.

### Gene repositioning does not correlate with gene activity

Finally, we asked if the changes in position of hypoxia-responsive genes correlate with the extent of their activation. We first assessed the 21 genes which showed statistical differences between normoxia and hypoxia in the radial distance analysis. Of those, 16 genes were up-regulated, whereas 5 genes were down-regulated upon hypoxic treatment for 48 h (Figure 4A). Up- and down- regulated gene groups contained both internal and external-shifted genes, and no clear relationship between radial position and expression profile was observed (Figure 4A).

**Figure 4:**
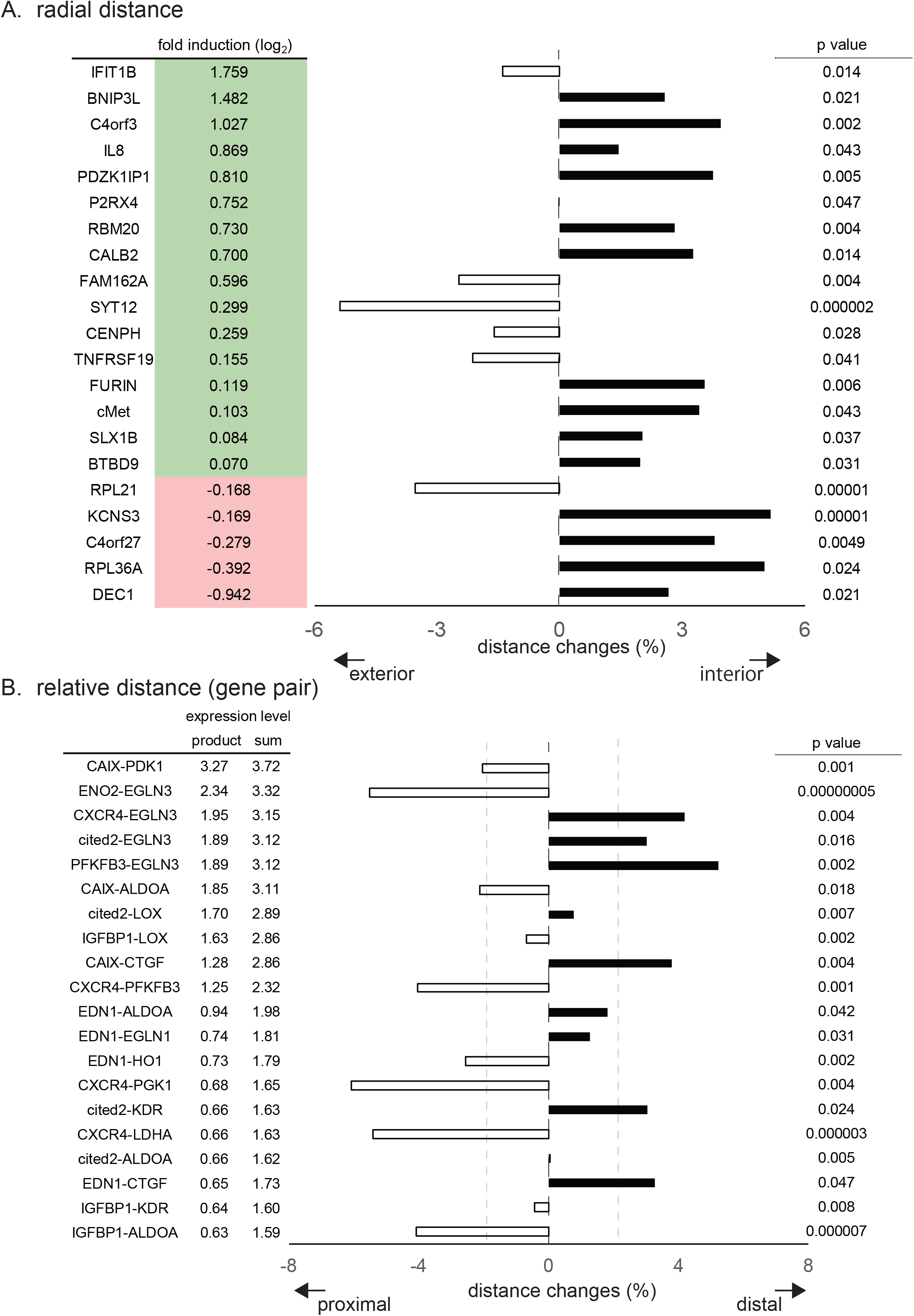
Expression profile of genes that change radial and relative distances under hypoxic conditions. mRNA-seq analyses were performed on MB231 cells treated with or without hypoxic condition for 48 h (n=2). Correlation of gene expression, and radial (A) or relative (B) distance was analyzed. Expression profiles of genes whose distance changed significantly in the radial distance analysis is shown as a fold induction (log2) upon hypoxic treatment (A, left). Green: up-regulated genes, red: down-regulated genes. Expression level of the two genes in the pair was expressed as the product and sum of the induction level of two genes in the relative distance analysis (B, left).

To assess the relationship of gene expression and relative positioning of gene pairs, the expression level of the two genes in the pair was expressed as the sum or product of the induction level of two genes. i.e., a larger product indicates larger change in the expression of corresponding two genes upon hypoxia; induction or repression of both genes will yield a positive value, opposite responses a negative value. The top 20 gene pairs which showed the largest change in the expression level were assessed. There were 10 gene pairs whose distance became closer and 10 pairs which separated, indicating that changes in the distance and expression level have no apparent relationship (Figure 4B; P < 0.05). An example is *ZEB1* on chromosome 10 and *VEGF* on chromosome 6 which upon hypoxic treatment moved apart and also showed opposite expression behavior with *ZEB1* down-regulated and *VEGF* up-regulated (Figure 3B, Supplemental Table 5, top line). However, this is not a general pattern, since *ZEB1* became more proximal to another up-regulated gene, *EGLN1* on chromosome 1 (Figure 3C, Supplemental Table 5, bottom line).

Taken together, these results suggest that changes in gene position are not strictly linked to gene expression status and they further point to lack of clustering of co-regulated genes.

## Discussion

Genes occupy non-random positions in the 3D space of the cell nucleus but the relationship between gene position and gene activity has remained unclear (Takizawa *et al*., 2008b). While gene position has been linked to gene expression levels mostly in studies of individual or small sets of genes (Zink *et al*., 2004; Takizawa *et al*., 2008b; Meaburn *et al*., 2009; Leshner *et al*., 2016; Forsberg *et al*., 2019), no large scale analysis relating gene location to activity has been reported. In this study, we used hypoxia as a model system and high-throughput imaging to map the location of nearly 100 genes and to analyze their spatial position in the nucleus and their repositioning behavior upon hypoxic stimulation (Schito and Semenza, 2016). Analysis was performed for the target genes of hypoxia-activated transcription factors HIF and CREB, and their nuclear positions were assessed by radial and relative distance analysis.

We find that in a set of 104 genes, all of them occupy preferred, non-random radial positions, yet all genes showed significant variability in their position in individual cells (Supplemental Table 2). These observations are in agreement with mapping of individual or small sets of genes over the years (Zink *et al*., 2004; Takizawa *et al*., 2008a; Takizawa *et al*., 2008b; Meaburn *et al*., 2009; Leshner *et al*., 2016; Forsberg *et al*., 2019) and support the notion that most genes occupy a preferred, but probabilistic 3D distribution in the cell nucleus.

Upon hypoxic treatment, only a subset of genes (∼ 20%) changed their radial position. Radial distance analysis demonstrated that 2 out of 35 genes (5.7%) in the HT group, but 18 out of 64 genes (28.1%) in the ICT group altered their positions with statistical significance upon hypoxic treatment, indicating that CREB target genes are more likely to reposition than HT genes. This difference may indicate that repositioning of genes depends on the signaling pathway which is activated, or they may reflect the fact that CREB is getting activated, whereas HIF-1α is becoming inactivated, at 48 h of hypoxic treatment. Relative gene position analysis was only performed for the HT group in the present study, and it remains unknown for the ICT or CDT groups. Considering the greater changes of radial position for the ICT and CDT group genes, it is possible that these groups also exhibit changes in their relative gene position. Importantly, two genes from the HT group, cMet and DEC1, which repositioned in the radial distance analysis, were also included in 6 of the gene pairs which repositioned in the relative distance analysis. These results represent, to our knowledge, the first analysis of the behavior of a large set of co-regulated genes with regards to their nuclear locations and they suggest that neither gene activation nor inactivation is a major determinant of radial gene location.

An attractive idea in the field has been that co-regulated genes cluster in 3D space, for example, via association with transcription factories or nuclear bodies which contain the necessary transcription factors for their activation. Correlations between gene expression and gene position, including the suggestion of clustering of genes, have been reported in several systems such as B lymphocyte development, ES cell differentiation and glial differentiation (Kosak *et al*., 2002; Williams *et al*., 2006; Osborne *et al*., 2007). On the other hand, there are also reports indicating that no clear correlation exists between gene position and expression (Meaburn and Misteli, 2008). Analysis of the pairwise distances of 159 combinations of 35 HIF-target genes showed limited changes in relative positioning, suggesting that coordinated relocation or clustering of co-regulated genes is not a pervasive phenomenon.

The 3D location of genes has been used to monitor cellular states, including to distinguish normal from cancer tissues (Meaburn, 2016). For example, *SP100* and *TGFB3* localize more peripherally in prostate cancer tissues compared to normal prostate (Meaburn and Misteli, 2019). However, as observed in our study, in earlier analyses the location of the repositioning genes was unrelated to their activity status (Meaburn and Misteli, 2008; Therizols *et al*., 2014; Shachar *et al*., 2015). Similar to breast and prostate cancer, and despite the seemingly limited relationship of gene activity with spatial position, it may be attractive to use gene positioning as a marker for hypoxia in tissues. This approach may overcome the notorious challenge posed by the highly unstable nature of HIF to assess the hypoxic state of tissues. The genes identified here which reposition may be promising candidates to do so, regardless of their activation status. This approach could for example be applied to assess the degree of hypoxia in tumor specimens in response to the HIF-2α inhibitor Belzutifan which is in clinical trials for renal cancers (Courtney *et al*., 2018, Choueiri *et al*., 2021) and approved recently by FDA.

Taken together, this study reports the nuclear positioning behavior of the largest set of genes in response to stimulation. These results based on the largest gene set analyzed to date, strongly support the lack of strict relationship of radial position and gene activity and our findings highlight the heterogeneous response of individual genes upon changes in gene expression.

## Materials and Methods

### Cell culture

MDA-MB231cells were obtained from ATCC (Manassas, VA) and cultured in Dulbecco’s modified Eagle’s medium (high-glucose) (Invitrogen, Carlsbad, USA) containing 10% fetal bovine serum (FBS) and antibiotics.

### Hypoxic treatment

Cells were treated under hypoxic conditions (1% O2 and 5% CO2, balanced with N2) in a hypoxia chamber (Billups-Rothenberg, Inc, Del Mar, CA for FISH experiments) or in a hypoxia workstation (Hirasawa Works, Tokyo, Japan for mRNA-seq analysis). An oxygen sensor was used to regulate the oxygen concentration inside the workstation, which was maintained at 1% throughout the experiment (MC-8G-S, Iijima Electrics, Gamagori, Japan).

### Oligonucleotide FISH probes

A pool of oligonucleotides for 104 hypoxia-inducible genes (an average of 819 oligonucleotides/gene within a 100 kb region centered around the gene) was synthesized (Twist Bioscience, South San Francisco, CA). Briefly, oligonucleotide probes contained a genomic sequence of the gene which is connected with a 32-bp barcode sequence that hybridizes to secondary-probes. The oligo pool was amplified by PCR, and used as a template for T7 primer-based *in vitro* transcription to synthesize primary probes (Beliveau *et al*., 2014). Fluorescently labeled secondary readout probes (Eurofin Genomics, Louisville, KY) which hybridize with specific barcode sequences on the primary probes, were used for detection (Beliveau *et al*., 2012).

### Oligo paint-based High-Throughput FISH in 384-Well Plates

For high-throughput FISH, cells were plated in 384-well CellCarrier plates (Perkin-Elmer) at a concentration of 2,000 cells/well. After normoxic or hypoxic culture for up to 48 h, cells were fixed in 4% paraformaldehyde in PBS for 15 min. After two washes with PBS, cells were permeabilized in 0.5% Triton X-100, 0.5% Saponin/PBS for 20 min at room temperature (RT) and then incubated in 0.1 N HCl for 15 min at RT. Cells were kept in 50% formamide/2x SSC for at least 30 min at RT. A probe mix containing 60 ng of each library probe and fluorescently labeled readout probe and 40 μg human COT1 DNA (Invitrogen) in 1.1 ml of hybridization buffer (20% dextran sulfate, 50% formamide, 2x SSC, 1x Denhartd’s solution) was used. 15 ul probe mix was added to the corresponding wells of 384-well plate prepared using a JANUS liquid handler workstation (PerkinElmer, Waltham, MA). Probes were denatured at 85°C for 7 min, and the plate was incubated at 37°C overnight (16 h) for hybridization. After incubation, excess probe was washed off three times with 100 ul 2x SSC, 2x SSC at 42°C, and 2x SSC at 60°C for 5 min each. Cells were stained with DAPI in PBS (5 ng/ml) before imaging.

### Image Acquisition and Analysis

Cells were imaged in 384-well plates on a CV7000 confocal high-throughput imaging system (Yokogawa Inc., Tokyo, Japan) using 4 solid state laser lines (405, 488, 561, 640 nm) for excitation, a 405/488/561/640 nm excitation dichroic mirror, a 40X air objective lens, a 568 emission dichroic mirror, and 2 sCMOS cameras (Andor), matched with appropriate emission bandpass filters (445/45, 525/50, 600/37, and 676/29 nm). Camera pixel binning was set to 2X2, for a resulting pixel size of 325 nm. Images were acquired in 4 channels as z-stacks of a total of 4 microns with images acquired at 1.0 μm steps. At least 6 randomly sampled fields were imaged per well. All image analysis steps were performed using Konstanz Information Miner (KNIME) software as described (Gudla et al., 2017). First, images from the same field of view and channel were maximally projected. Then, nuclei were segmented using the DAPI channel. The resulting nucleus ROI was used as the search region for the FISH spot detection algorithm in the Alexa 488, ATTO 550, and Cy5 channels, respectively. The normalized radial distance of each nucleus ROI pixel was then measured by dividing each absolute radial distance value by the per-cell maximum radial distance value. The nucleus border assumes a normalized value of 0, whereas the nucleus center has a normalized value of 1. The normalized absolute radial position of the FISH signal was calculated at the spot center pixel (Gudla *et al*., 2017).

### Statistical Analysis

For all experiments we conducted two to six times of experiments under the experimental condition of hypoxia or normoxia (biological replicates) and measured the nuclear radial position of genes (median of 4409 and 4600 cells imaged in normoxia and hypoxia per biological replicate, respectively) and the relative distance between the two genes (median of 1942 and 1687 cells imaged in normoxia and hypoxia per biological replicate, respectively) in each experiment.

Genes or gene pairs whose radial or relative distance were subjected to Wilcoxon rank sum test, and the genes which were statistically different (p value smaller than a Bonferroni significance threshold) between biological replicates conducted under the same experimental conditions were excluded from the analysis for evaluating the association between the experimental conditions and the radial and relative distances. The Wilcoxon rank-sum test was used to compare the radial and relative distance between biological replicates. P-values of less than a Bonferroni significance threshold determined by the number of pairwise comparisons between biological replicates for the radial or relative distance were considered to indicate statistically difference. For each gene and gene pair whose radial or relative distances were not statistically different between biological replicates as a result of the above analysis, we performed an experiment-level meta-analysis to evaluate the association between the experimental conditions and the nuclear radial and relative distances. Specifically, in each set of experiments performed under hypoxia and normoxia, we first calculated the ratio of means of radial and relative distances (hypoxia / normoxia) and its variance, and the p value on the Wilcoxon rank sum test. Next, we calculated the weighted ratio of means of radial and relative distances and corresponding 95% confidence interval using the inverse variance-weighted average method (Friedrich et al. 2011). Additionally, we used Fisher’s method (Fisher 1932) to combine multiple raw p-values from independent experiments. P values were adjusted for multiple comparisons with the use of the method of Benjamini and Hochberg (Benjamini and Hochberg, 1995) to control the false discovery rate at the 0.05 level. All statistical analyses were performed using SAS version 9.4 (SAS Institute 193 Inc., Cary, NC, USA). Statistical significance was defined as corrected p<0.05.

### mRNA-seq analysis

The mRNA expression profiles of MB231 cells treated with normoxia (21% O2) or hypoxia (1% O2) for 48 h were analyzed. RNA sequence library preparation, sequencing, mapping, and gene expression analysis were performed by DNAFORM (Yokohama, Japan). Qualities of total RNA were assessed by Bioanalyzer (Agilent, Santa Clara, CA). After poly (A) + RNA enrichment by NEBNext Poly(A) mRNA Magnetic Isolation Module (New England BioLabs, Ipswich, MA), RNA-seq library was prepared using the SMARTer Stranded Total RNA Sample Prep kit - HI Mammalian (Takara Bio Inc, Shiga, Japan) following the manufacturer’s instructions. Then first-strand synthesis was performed using an N6 primer. Illumina specific indexed libraries were amplified by PCR. The libraries were sequenced on a NextSeq 500 sequencer (Illumina, San Diego, CA) to generate 50 nt and 25 nt paired end reads. Obtained reads were mapped to the human GRCh38.p10 genome using STAR (version 2.7.2b). Reads on annotated genes were counted using featureCounts (version 1.6.1). FPKM values were calculated from mapped reads by normalizing to total counts and transcript (Supplemental Table 6).

Differentially expressed genes were detected using the DESeq2 package (version 1.20.0). Sequence results were deposited in the DDBJ Sequence Read Archive database (accession number: SSUB018734).

### Lactate assay

MB231 cells were cultured under normoxic or hypoxic conditions for 48 h, and lactate levels in the medium were measured using a Lactate Assay Kit as per the manufacturer’s instructions (Biovision, Milpitas, CA).

### Immunofluorescence

Cells were fixed with 25μl of 4% paraformaldehyde per well (10 min), permeabilized for 10 min (PBS/0.5% triton-X 100), and washed once with PBS/0.05% Tween-20. Then, cells were incubated for 1 h with primary antibodies (anti-HIF-1α, 1:100, BD Bioscience #610958, anti-phospho CREB, 1:200, CST #9198S) diluted in blocking buffer (PBS/0.05% Tween 20/ 5% bovine serum albumin). After three washes in wash buffer (PBS/0.5% Tween-20), cells were incubated for 1 h with secondary antibodies (anti-Mouse IgG 488, 1:1000 Invitrogen, #A11029; anti-rabbit IgG 488, 1:1000, Invitrogen, #A11034), followed by DAPI staining (5 μg/ml).

## Acknowledgment

We thank Gianluca Pegoraro, Reddy Gudla and Laurent Ozbun for help with high-throughput imaging and image analysis performed in the NCI High-throughput Imaging Facility T.M. was supported by funding from the Intramural Research Program of the NIH, the NCI Center for Cancer Research. K.N. was supported by the Princess Takamatsu Cancer Research Fund and a Grant-in-Aid for Scientific Research (JSPS KAKENHI Grant Number 15KK0298,18KK0233).

---

**Supplemental Figure 1.**
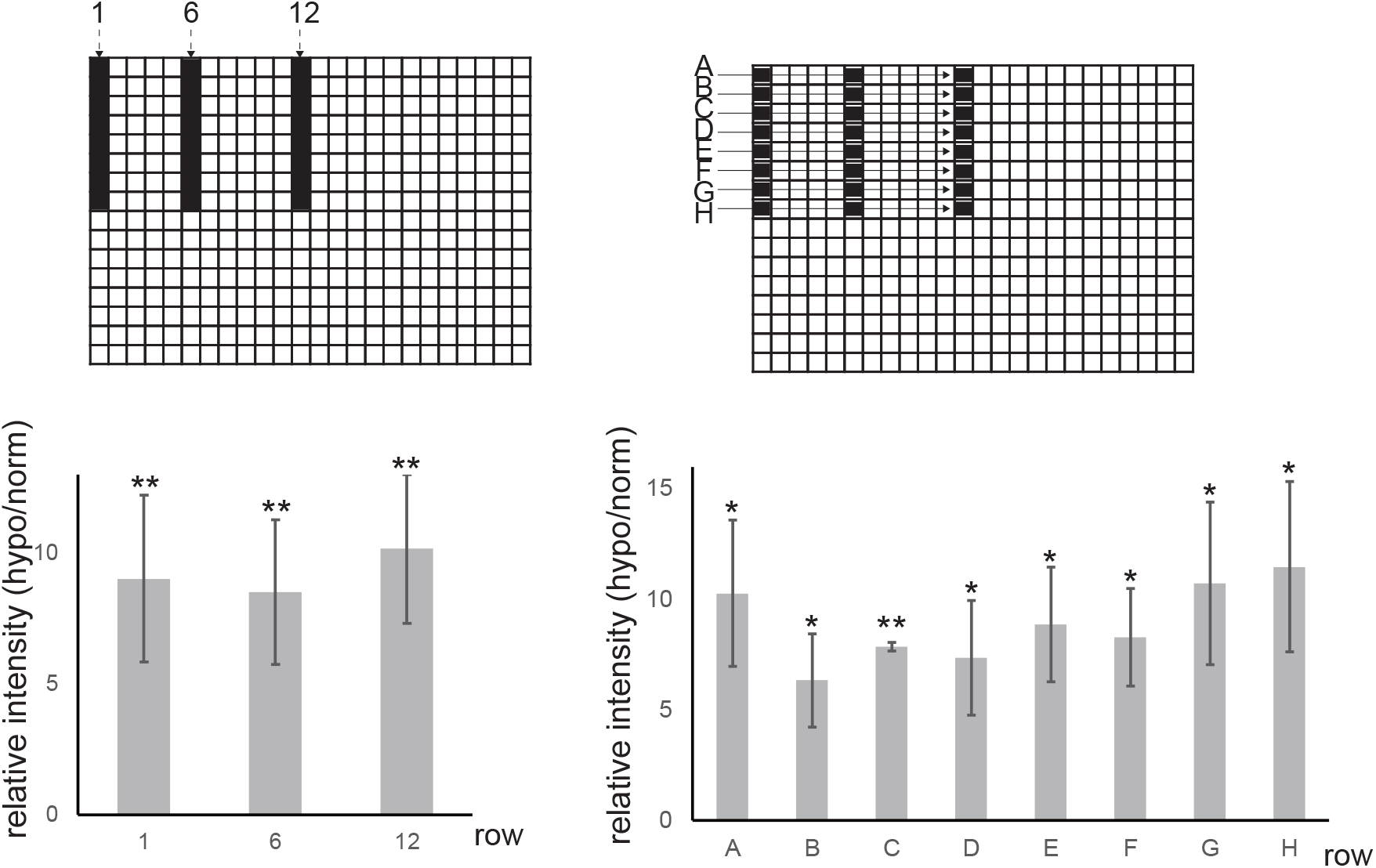
Brightness of stained-HIF1α image was measured and averaged by row; 1, 6, 12 (n=8), A-H (n=3). Error bars indicate SD. Significance was analyzed by t-test (*p<0.05, **p<0.02).

**Supplemental Figure 2.**
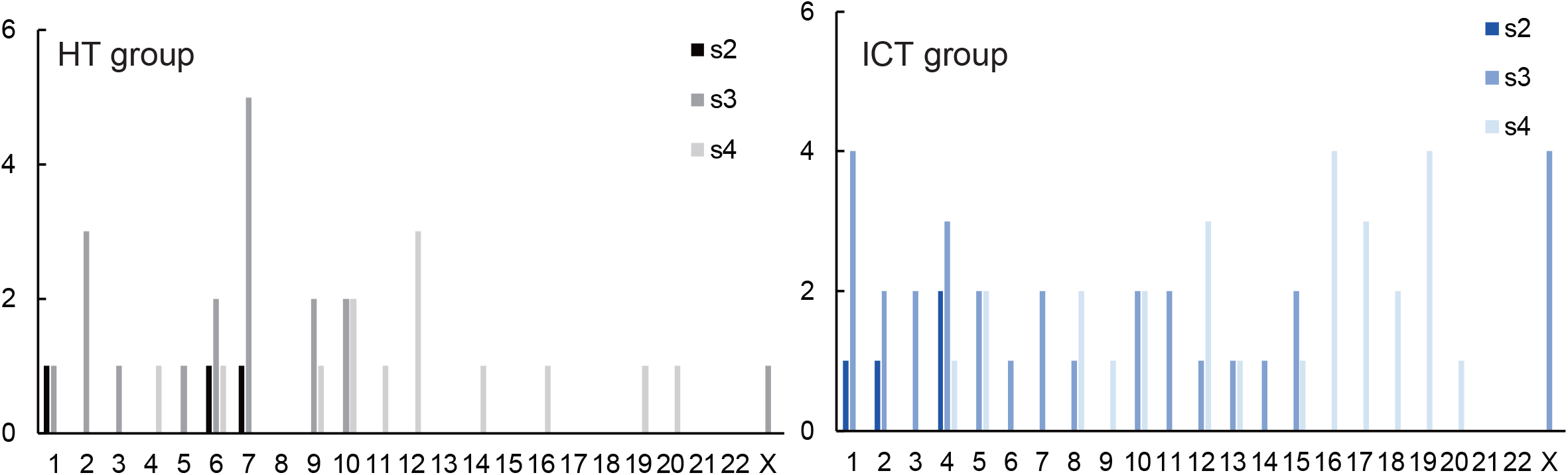
Relationship between radial distance and chromosome location. Genes of the HT and ICT group are classified based on their chromosome location and plotted in relation to the radial distance (equi-area analysis shell number(s)2, 3 or 4). Gene number was counted in the integrated data of two experiments. Black/grey: HT group, blue/light blue: ICT group.

**Supplemental Table 1.**
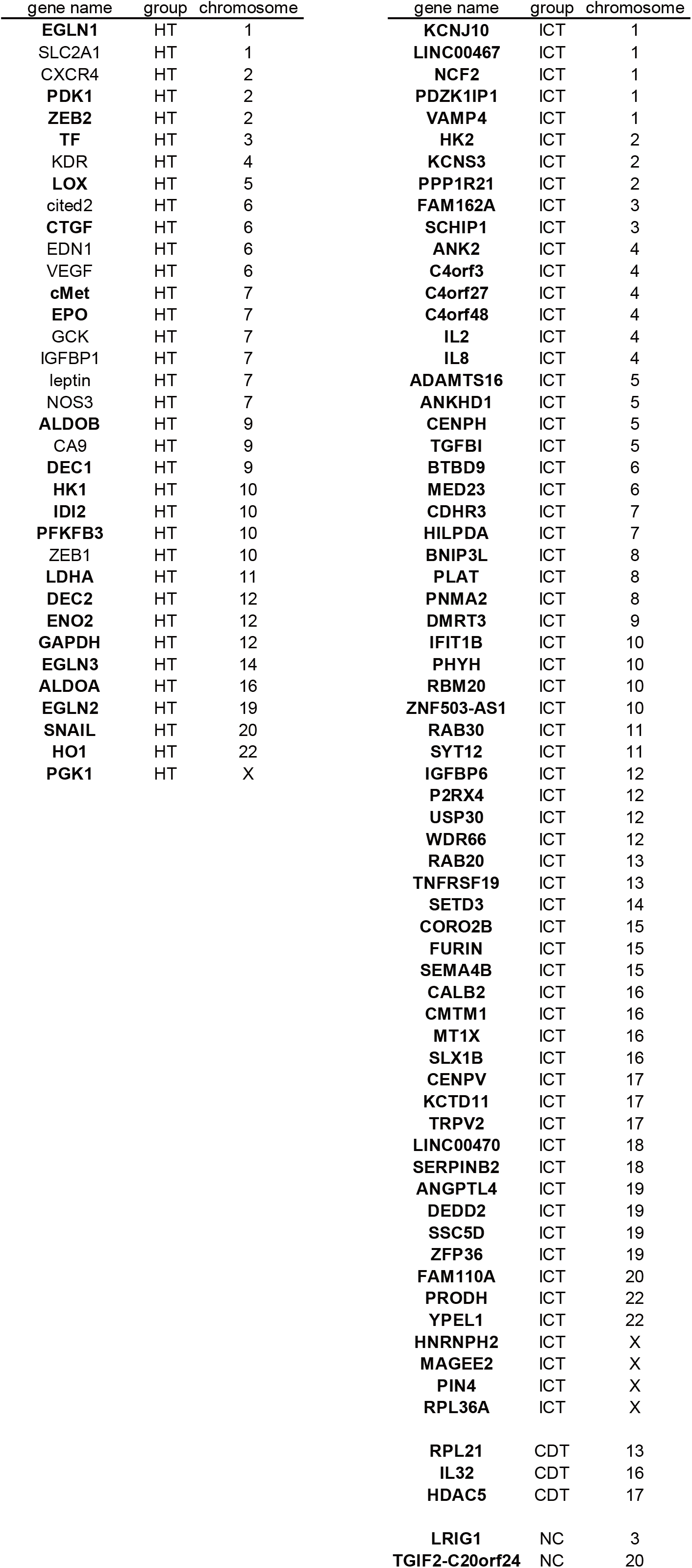
Genes analyzed in the study

**Supplemental Table 2.**
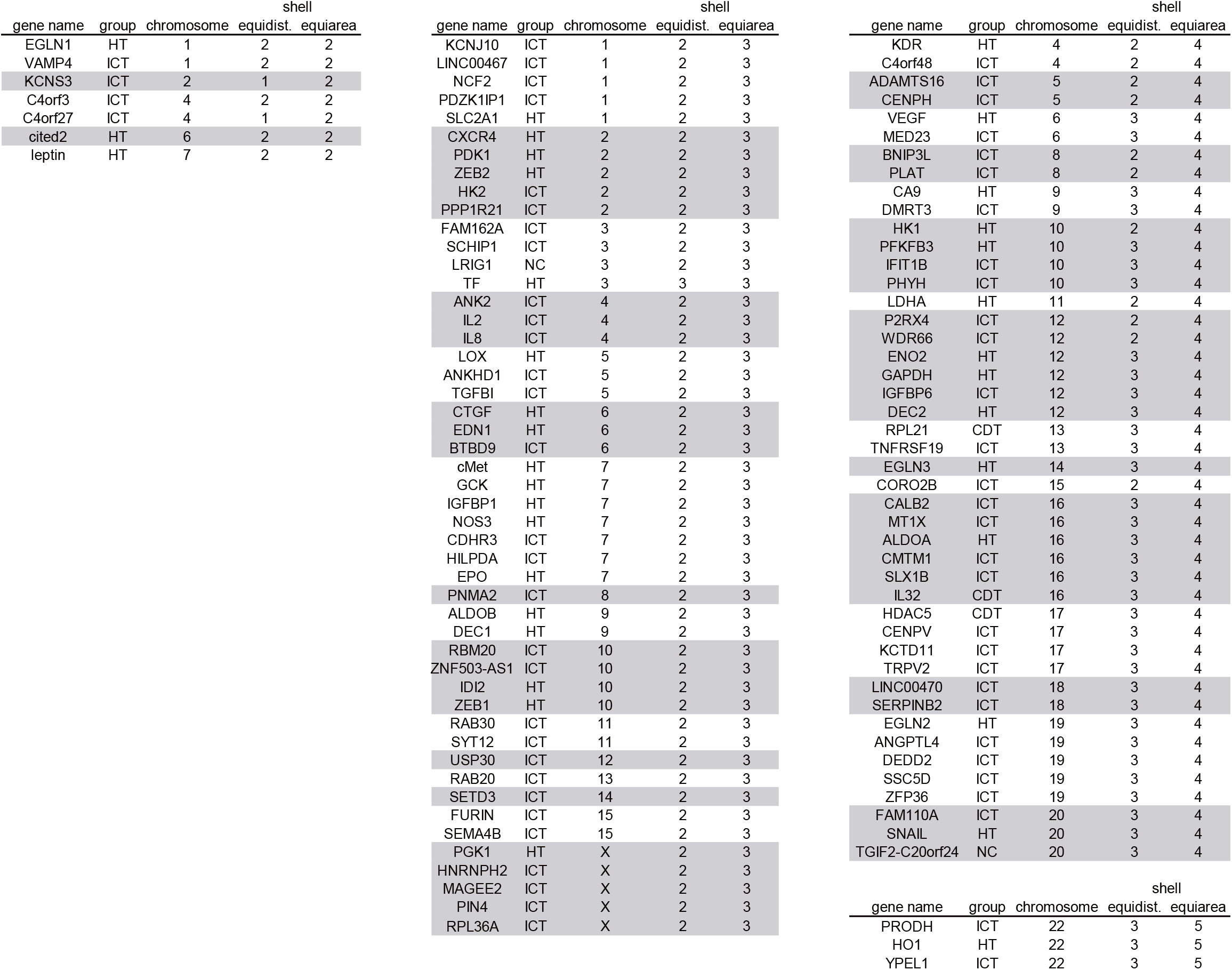
Chromosome locations of genes and their shell positions in the nucleus

**Supplemental Table 3.** Radial distance of genes comparing normoxic and hypoxic conditions blue: genes shifted toward center of nuclei, yellow: those shifted toward periphery green: genes up-regulated in hypoxia, red: genes down-regulated in hypoxia

| gene name | chromosome | p value (FDR) | ratio of mean (hypo/norm) | 95% confidence interval |  | fold induction (log<sub>2</sub>) |
| --- | --- | --- | --- | --- | --- | --- |
|  |  |  |  | lower bound | upper bound | hypo/norm |
| KCNS3 | 2 | 0.00001 | 1.052 | 1.027 | 1.077 | -0.169 |
| RPL36A | X | 0.024 | 1.050 | 1.016 | 1.086 | -0.392 |
| C4orf3 | 4 | 0.002 | 1.039 | 1.017 | 1.062 | 1.027 |
| PDZK1IP1 | 1 | 0.005 | 1.038 | 1.017 | 1.059 | 0.810 |
| C4orf27 | 4 | 0.0049 | 1.0379 | 1.01 | 1.06 | -0.279 |
| FURIN | 15 | 0.006 | 1.036 | 1.011 | 1.061 | 0.119 |
| cMet | 7 | 0.043 | 1.034 | 1.011 | 1.057 | 0.103 |
| CALB2 | 16 | 0.014 | 1.033 | 1.012 | 1.054 | 0.700 |
| RBM20 | 10 | 0.004 | 1.028 | 1.009 | 1.048 | 0.730 |
| DEC1 | 9 | 0.021 | 1.027 | 1.009 | 1.045 | -0.942 |
| BNIP3L | 8 | 0.021 | 1.026 | 1.006 | 1.045 | 1.482 |
| SLX1B | 16 | 0.037 | 1.020 | 1.006 | 1.035 | 0.084 |
| BTBD9 | 6 | 0.031 | 1.020 | 1.005 | 1.034 | 0.070 |
| IL8 | 4 | 0.043 | 1.014 | 0.998 | 1.031 | 0.869 |
| P2RX4 | 12 | 0.047 | 1.000 | 0.986 | 1.014 | 0.752 |
| IFIT1B | 10 | 0.014 | 0.986 | 0.974 | 0.998 | 1.759 |
| CENPH | 5 | 0.028 | 0.984 | 0.969 | 0.999 | 0.259 |
| TNFRSF19 | 13 | 0.0405 | 0.979 | 0.96 | 1.00 | 0.155 |
| FAM162A | 3 | 0.004 | 0.975 | 0.958 | 0.993 | 0.596 |
| RPL21 | 13 | 0.00001 | 0.965 | 0.948 | 0.981 | -0.168 |
| SYT12 | 11 | 0.000002 | 0.946 | 0.925 | 0.968 | 0.299 |

**Supplemental Table 4.** List of gene pairs analyzed for relative distances

|  |  |  |  |
| --- | --- | --- | --- |
| ZEB1-VEGF | CXCR4-EGLN2 | CAIX-LDHA | CXCR4-LDHA |
| IDI2-EGLN3 | leptin-EGLN3 | CAIX-PGK1 | DEC1-GAPDH |
| CAIX-EGLN3 | ENO2-PGK1 | DEC1-DEC2 | ENO2-EGLN3 |
| EDN1-HK1 | ZEB2-LOX | IDI2-cMet | NOS3-ALDOB |
| cited2-EGLN3 | cited2-ALDOA | CXCR4-LOX | CXCR4-PGK1 |
| CTGF-HK1 | CTGF-EPO | IGFBP1-DEC2 | ZEB1-LOX |
| CAIX-EPO | ZEB2-PGK1 | IDI2-DEC2 | DEC1-LDHA |
| EDN1-cMet | cited2-HO1 | TF-PGK1 | ZEB1-HK1 |
| CXCR4-EGLN3 | TF-EGLN2 | ZEB1-GAPDH | ZEB1-EGLN1 |
| EDN1-GAPDH | ZEB1-PGK1 | TF-HK1 |  |
| IDI2-ALDOA | ENO2-HK1 | CXCR4-GAPDH |  |
| CAIX-CTGF | CAIX-GCK | leptin-PFKFB3 |  |
| CAIX-VEGF | NOS3-VEGF | leptin-GAPDH |  |
| EDN1-CTGF | SNAIL-LDHA | NOS3-GAPDH |  |
| EDN1-PGK1 | PFKFB3-EGLN2 | ZEB1-KDR |  |
| cited2-KDR | DEC1-EGLN2 | PFKFB3-DEC2 |  |
| EDN1-EGLN3 | leptin-ALDOA | SNAIL-ALDOB |  |
| PFKFB3-EGLN3 | DEC1-HO1 | cited2-PGK1 |  |
| CTGF-cMet | leptin-VEGF | leptin-EGLN2 |  |
| CAIX-GAPDH | ZEB1-LDHA | SNAIL-PGK1 |  |
| cited2-DEC2 | CTGF-DEC2 | leptin-HK1 |  |
| IGFBP1-VEGF | IGFBP1-KDR | CAIX-PDK1 |  |
| CAIX-EGLN2 | IGFBP1-HK1 | CTGF-ALDOB |  |
| EDN1-PFKFB3 | ZEB1-PDK1 | CAIX-ALDOA |  |
| SLC2A1-HK1 | SLC2A1-ALDOB | DEC1-KDR |  |
| EDN1-EPO | ZEB1-ALDOA | leptin-KDR |  |
| ZEB2-DEC2 | EDN1-GCK | leptin-ALDOB |  |
| EDN1-ALDOA | NOS3-KDR | ZEB2-cMet |  |
| EDN1-KDR | IGFBP1-LOX | CXCR4-HO1 |  |
| cited2-cMet | DEC1-HK1 | CXCR4-HK1 |  |
| CAIX-HK1 | CAIX-HO1 | CXCR4-CTGF |  |
| DEC1-LOX | ZEB1-EGLN3 | EDN1-HO1 |  |
| ZEB1-EGLN2 | CXCR4-VEGF | CXCR4-DEC2 |  |
| EDN1-EGLN1 | PFKFB3-GCK | IDI2-HO1 |  |
| IDI2-EGLN2 | NOS3-HO1 | IDI2-GCK |  |
| IDI2-PGK1 | CXCR4-ALDOA | ZEB1-GCK |  |
| CTGF-EGLN2 | SNAIL-GAPDH | leptin-DEC2 |  |
| NOS3-HK1 | IGFBP1-ALDOB | CXCR4-KDR |  |
| DEC1-ALDOA | ZEB1-DEC2 | EDN1-EGLN2 |  |
| EDN1-SNAIL | SLC2A1-KDR | DEC1-cMet |  |
| TF-EGLN3 | IDI2-LOX | TF-ALDOA |  |
| CAIX-cMet | EDN1-DEC2 | SNAIL-GCK |  |
| cited2-LOX | cited2-HK1 | SNAIL-KDR |  |
| cited2-GCK | ZEB1-cMet | SNAIL-DEC2 |  |
| SNAIL-HK1 | ZEB1-ALDOB | EDN1-ALDOB |  |
| TF-HO1 | ZEB2-HK1 | leptin-CTGF |  |
| ENO2-EGLN2 | ZEB2-ALDOB | CXCR4-PFKFB3 |  |
| ZEB1-HO1 | CAIX-KDR | IGFBP1-ALDOA |  |
| ZEB1-EPO | SNAIL-EGLN2 | DEC1-PGK1 |  |
| CAIX-PFKFB3 | leptin-LDHA | CXCR4-EPO |  |

**Supplemental Table 5.**
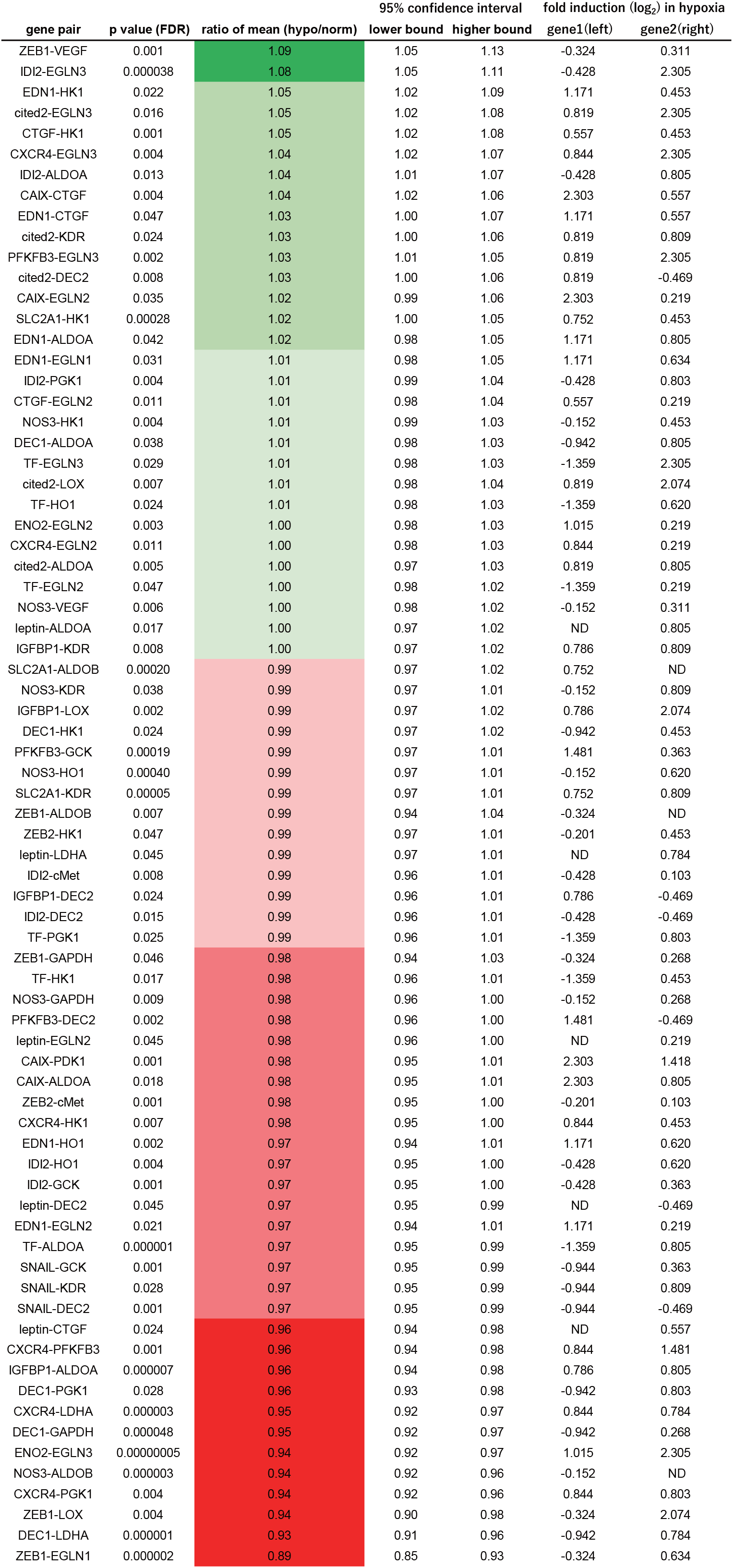
**Gene pairs whose distribution of relative distance changed statistically significant under hypoxic condition** green: gene pairs whose distance became farther, red: closer

**Supplemental Table 6.** Number of sequence reads

| **sample** | **total input reads** | **uniquely mapped reads** | **multi-mapped reads** |
| --- | --- | --- | --- |
| 0 h-rep1 | 52,127,618 | 32,916,779 | 6,300,162 |
| 0 h-rep2 | 46,831,702 | 30,227,426 | 6,266,476 |
| 48 h-rep1 | 45,543,343 | 27,575,569 | 5,084,523 |
| 48 h-rep2 | 58,840,386 | 32,857,007 | 6,304,537 |

